# Fibro-adipogenic Progenitor and Macrophage Remodeling of the Aging Skeletal Muscle Niche During Exercise-Induced Hypertrophy

**DOI:** 10.64898/2026.09.25.754304

**Authors:** Jakob Wang, Jonas Brorson, Søren Kuhdal, Jesper Just, Frederik Forsberg Thybo, Mette Bjerre, Frank Vincenzo de Paoli, Niels Jessen, Kristian Vissing, Jean Farup

**Affiliations:** Dept. of Biomedicine, Aarhus University, Aarhus, Denmark; Steno Diabetes Center Aarhus, Aarhus University Hospital, Aarhus, Denmark; Nuclear Medicine and PET Center, Aarhus University Hospital, Aarhus, Denmark; Department of Molecular Medicine, Aarhus University Hospital, Aarhus, Denmar; Department of Clinical Medicine, Aarhus University, Aarhus, Denmark; Dept. of Thoracic Surgery, Aarhus University Hospital, Aarhus Denmar; Dept. of Public Health, Aarhus University, Aarhus, Denmark

**Keywords:** aging, strength training, mesenchymal stem cells, inflammation, exercise adaptation

## Abstract

Fibro-adipogenic progenitors (FAPs) have emerged as central regulators of the skeletal muscle homeostasis and the muscle microenvironment. However, the role of FAPs in the context of exercise-induced muscle hypertrophy remains largely unexplored. Here, we utilized six weeks of blood flow restricted resistance exercise (BFRRE) to study cellular adaptations within the muscle microenvironment accompanying muscle hypertrophy in healthy older individuals. Using flow cytometry, we characterized global changes of key cell populations within the skeletal muscle microenvironment, including FAPs, muscle stem cells (MuSCs), and immune cells. BFRRE induced significant enlargement of both FAPs and MuSCs, consistent with cellular adaptation to exercise. Notably, exercise shifted the FAP pool toward an increased predominance of the CD90^high^ FAP phenotype, without altering total FAP abundance. In addition to matrix and collagen-related genes, transcriptional analysis of genes associated with secretory proteins revealed enrichment of pro-myogenic factors in CD90^high^ versus CD90^low^ FAPs. Conditioned media experiments of freshly isolated FAPs demonstrated that CD90^high^ FAPs promote myotube growth *in vitro* compared to CD90^low^ counterparts, suggesting that this phenotypic shift may facilitate muscle hypertrophy. In parallel, BFRRE increased the proportion of pro-inflammatory (CD11c+) macrophages within the skeletal muscle niche, highlighting a dynamic immune response during adaptation. Finally, we show identified possible link between pro-inflammatory macrophages and FAPs, as TNFα markedly reduced the proliferation of human primary FAPs *ex vivo*, suggesting that macrophage-derived signals may attenuate excessive FAP expansion during tissue remodelling. Together, these findings provide new insight into how the remodelling of the cellular niche may support muscle hypertrophy in response to exercise. The coordinated expansion and phenotypic remodeling of FAP and immune cell populations may represent an important mechanism through which exercise supports hypertrophy in older individuals.

**Key Points:**

- The role of FAPs in human exercise-induced hypertrophy remains poorly understood.
- Exercise-induced hypertrophy increased FAP size and the proportion of CD90^high^ FAPs.
- CD90^high^ FAPs expressed growth-promoting ligands and enhanced myotube growth through secreted factors compared to CD90^low^ FAPs.
- CD11c⁺ macrophage expansion correlated with CD90^high^ FAP and MuSC responses, while TNF-α inhibited FAP proliferation ex vivo.
- The results highlight remodelling of FAP subpopulations as a potential contributor to exercise-induced hypertrophy of aging skeletal muscle.

## Introduction

Skeletal muscle is essential for locomotion, metabolic homeostasis, and overall health, accounting for approximately 40% of total body mass in healthy adults [1, 2]. Aging is accompanied by progressive declines in muscle mass and quality, characterized by myofiber atrophy, fibro-fatty infiltration, impaired regenerative capacity, and reduced contractile function [2, 3]. These changes contribute substantially to frailty, disability, and mortality in older adults. Physical activity, and in particular resistance-based exercise remain one of the most effective strategies to counteract age-related deterioration of skeletal muscle.

The mechanisms orchestrating skeletal muscle adaptation to exercise are increasingly recognized to extend beyond the muscle fibers themselves and involve coordinated interactions among multiple cell populations within the muscle microenvironment [4, 5]. Among these, fibro-adipogenic progenitors (FAPs) have emerged as central regulators of skeletal muscle remodeling in health and disease [6–8]. These mesenchymal, stromal cells interact with muscle stem cells (MuSCs), immune cells, and the extracellular matrix, to regulate tissue repair through dynamic [9–13]. Following muscle injury, FAPs rapidly expand and promote a transient pro-regenerative niche through secretion trophic and chemotactic factors [10, 14, 15]. Meanwhile, timely macrophage-mediated termination of FAP activity is important to prevent excessive FAP accumulation, which can result in displacement of fat and connective tissue and subsequent muscle dysfunction [10]. Analogously, alterations in FAP function and their excessive accumulation have been implicated in the fibro-fatty degeneration and muscle wasting observed aging and chronic diseases [5, 15–19]. On the other hand, experimental depletion of FAPs results in a dramatic loss of muscle mass, function, and regenerative capacity in mice [5, 20, 21]. Collectively, these findings highlight FAPs as important regulators of muscle homeostasis, while emphasizing their context-dependent and pleiotropic role in governing muscle health and quality [22].

While the role of FAPs as drivers of pathological muscle remodeling is well established, their contribution to physiological muscle adaptation, remains poorly defined. Resistance exercise effectively counteracts manifestations of the aging skeletal muscle phenotype by promoting myofiber hypertrophy, enhancing regenerative capacity, and suppressing fibro-fatty remodeling [23–26]. Pre-clinical studies identifies FAPs amongst the most exercise-responsive cells throughout the organism [27], with increased FAP activity and paracrine signaling being linked extracellular matrix and niche remodeling [28, 29]. Recently, the exercise-responsiveness of skeletal muscle FAPs has been tied to mechanosensitive mechanisms, inferring that resistance exercise may represent a potent regulator of FAP content, composition, and function [30]. Oppositely, prolonged exercise has been shown to induce senescence of FAPs in mice [31]. None-the-less, evidence from human studies are lacking and it remains unknown whether distinct FAP subpopulations may contribute to exercise-induced muscle hypertrophy.

We have demonstrated that human skeletal muscle FAPs comprise distinct sub-populations with divergent functional properties, with CD90^high^ FAPs characterized by high metabolically activity and collagen production [32]. Notably, we also recently observed that muscle wasting in patients with cancer cachexia was associated with a relative reduction in CD90^high^ FAPs, and that FAPs isolated from the cachectic muscle promoted myotube atrophy *in vitro* compared with FAPs from healthy controls [17]. These findings raise the possibility that specific FAP sub-populations are important for the regulation of muscle mass in humans.

Here, we investigated whether hypertrophy-inducing exercise was associated with changes in the phenotype of the FAP compartment of human skeletal muscle. Using low-load blood flow restriction exercise (BFRRE), as an established model to promote exercise-induced hypertrophy [33], we hypothesized for increases FAP content to underlie the anabolic response to resistance exercise.

We found that exercise-induced hypertrophy was associated with markedly increased the proportion and size of CD90^high^ FAPs within the skeletal muscle. Through conditioned media experiments, we confirmed that CD90^high^ FAPs promotes myotube growth through secreted factors *in vitro*. Interestingly, the observed alterations of the FAP compartment were associated with increases in the proportion of pro-inflammatory macrophages within the muscle. Functionally, we identified a potential mechanistic link between FAPs and pro-inflammatory macrophages, as TNF-α markedly reduced the proliferation of human primary FAPs *ex vivo*. Collectively, these findings suggest that changes in specific subsets of FAPs can drive muscle growth in humans.

## Methods

### Study design and exercise intervention

For detailed information of the study design the reader is referred to the original study [33]. Briefly described, participants were randomized to either six weeks of BFRRE (three training sessions pr. week) or a corresponding non-exercise control period (CON). Each BFRRE training session comprised 5 minutes stationary bicycling followed by a standardized low-intensity warmup of the knee extensors. This was immediately followed four sets of knee extensions at 30% of 1RM, with all sets performed to volitional failure while BFR was applied at 50% of the arterial occlusion pressure. The study was conducted in accordance with the Declaration of Helsinki and approved by the Central Region Denmark Committee on Health Research Ethics (1-10-72-169-20) and registered at ClinicalTrail.gov (NCT04712955).

### Participants

Detailed information of the study population is provided in the original study [33]. A total of 23 participants (two females) were enrolled in the present study. Participants were characterized by a median age of 66 years (57-75 years), and within the normal range with regards to weight and height, equating to an average BMI 26.2±3.2 kg/m^2^. Exclusion criteria comprised any known chronic disease and regular intake of prescription medication (with exception of mild hypertensives and anti-depressants). Finally, from the time of inclusion, engagement in regular physical activity equivalent to >2 hours per week during the preceding year also excluded study participation. Block-randomization was stratified by sex and was conducted with the main investigator being blinded to block size. Samples for FACS analysis were missing from two participants from the original trial (one participant in each group), leaving the groups sizes n=11 (BFRRE) and n=10 (CON) in the present study.

### Biopsy sampling and tissue processing

Skeletal muscle biopsies were collected from the vastus lateralis using the Bergstrøm needle technique with manual suction [34]. All biopsies were collected in sterile conditions under local anesthesia (10 mL lidocaine, 10 mg/ml, Amgros I/S, Denmark). Pre-intervention biopsies were collected within a week of initiating the intervention, whereas post-biopsies were four days after the final exercise bout. Immediately after collection, biopsies were dissected free of visible fat and connective tissue and subsequently weighed, to enable normalized cell content assessments following FACS.

Tissue aliquots (appx. 170 mg) were buffered in C-tubes (cat. No 130-093-237, Miltenyi Biotech, Lund, Sweden) containing 7.5 mL of ice-cold wash buffer [Hams F10+ incl. glutamine and bicarbonate (cat. No. N6908, Sigma-Aldrich, Denmark) supplemented with 10% horse serum (cat. No. 26050088, Gibco, ThermoFisher Scientific, MA) and 1% pen/strep (cat. No. 15140122, Gibco, ThermoFisher Scientific, MA). Samples were kept on ice for no more than 4 hours prior to further processing.

A single cell suspension of mononuclear cells were achieved in accordance with previously published protocols from our group [35]. For tissue digestion, 700 U/ml collagenase II (cat. No 46D16552, Worthington, Lakewood, NJ) and 3.27 U/mL dispase II (cat. No. 04 942 078 011, Roche Diagnostics, Basel, Switzerland) was added to the C-tubes, immediately before mechanical disruption on the gentleMACS (cat. No 130-096-427, Miltenyi Biotech, Lund, Sweden) for 61 minutes using a skeletal muscle digestion program at 37 degrees (37C_mr_SMDK1) to achieve a single cell suspension. Following digestion, 10 ml of wash buffer was added C-tubes to dilute the enzymes and the single cell suspensions were transferred to 50 ml nunc tubes while passed through a 70-um cell strainer. C-tubes were washed twice with wash buffer to collect all remaining cells and transferred to the nunc tubes. Suspensions were centrifuged at 500 g for 5 minutes followed by removal of the supernatant. Finally, the cell pellet was resuspended in 1 ml of freezing buffer (StemMACS, cat. No 130-109-558) at stored at −80 degrees in cryovials until further preparation for FACS.

### Sample preparation and florescence activated cell sorting (FACS)

Samples were thawed in a 37C water bath until a small piece of ice was left in the cryovial (no more than 90 seconds) and immediately transferred to 12 ml of freshly prepared wash buffer before centrifuged at 500 g for 5 minutes. The supernatant was removed, and the cell pellet was resuspended in 125μ L wash buffer and incubated with the following antibodies; Anti-CD56-BV421 (5 μL/sample, cat. no. 562751, BD Biosciences, CA), Anti-CD82-PE-Vio770 (10 μL/sample, cat. no. 130-101-302, Miltenyi Biotec), Anti-CD34-APC (20 μL/sample, cat. no. 555824, BD Biosciences), Anti-CD31-PerCPvio 700 (4 μL/sample, cat. no. 700 130-110-673, Miltenyi Biotec), Anti-CD90-PE (3.6 μL/sample, cat. no. 12-0909-42, Invitrogen, ThermoFisher Scientific), Anti-CD45-VioBright FITC (2 μL/sample, cat. no. 130-114-567, Miltenyi Biotec), Anti-CD14-BV605 (5 μL/sample, cat. no. 564054, BD Biosciences) and Anti-CD11c-APCvio770 (2 μL/sample, cat. no. 130-113-585, Miltenyi Biotec). Incubation was performed at 5°C for 35 minutes, protected from light and with constant agitation. Following incubation, the suspension was diluted with 10 ml wash buffer and centrifuged at 500g for 5 minutes and subsequently the supernatant was removed. Finally, the cell pellet was resuspended in 300 µL wash buffer and transferred to FACS tubes passed through a 30 µm cell strainer, nunc tubes were washed twice in 300 μL wash buffer, that was also passed though the cell strainer. As so, the final volume for sorting was 900 µL pr. sample. Immediately before sorting 10 µL of proprium iodide (PI, cat. No. 556463, BD Bioscience) and absolute counting beads (Count-Bright, cat. No. C36959, ThermoFisher Scientific) were added to each sample to identify live cells and allow for assessment of cell counts normalized to mg tissue weight.

Flow cytometry and cell isolation were performed on FACS Aria-III (BD Bioscience) using 408-, 488-, 561-, and 633-nm lasers. A 100-µm nozzle at 20 psi was utilized to reduce the stress imposed on the cells and to prevent clogging. Live cells of interest were sorted into cooled 5 ml FACS collection tubes containing 500 µL of wash buffer. Gating strategy was set up in accordance with previous studies [32, 35, 36] and was optimized utilizing Fluorescence Minus One for all anti-bodies. Compensation beads (cat. No. 01-2222-41, eBioscience, ThermoFisher Scientific) were used to ensure bright single color stainings for compensation. Data acquisition was performed using FACS Diva software and subsequent analysis was performed using FlowJo Software, version 10.6.2. Gating strategies are depicted in *Supplementary Fig. 1*.

**Fig. 1.**
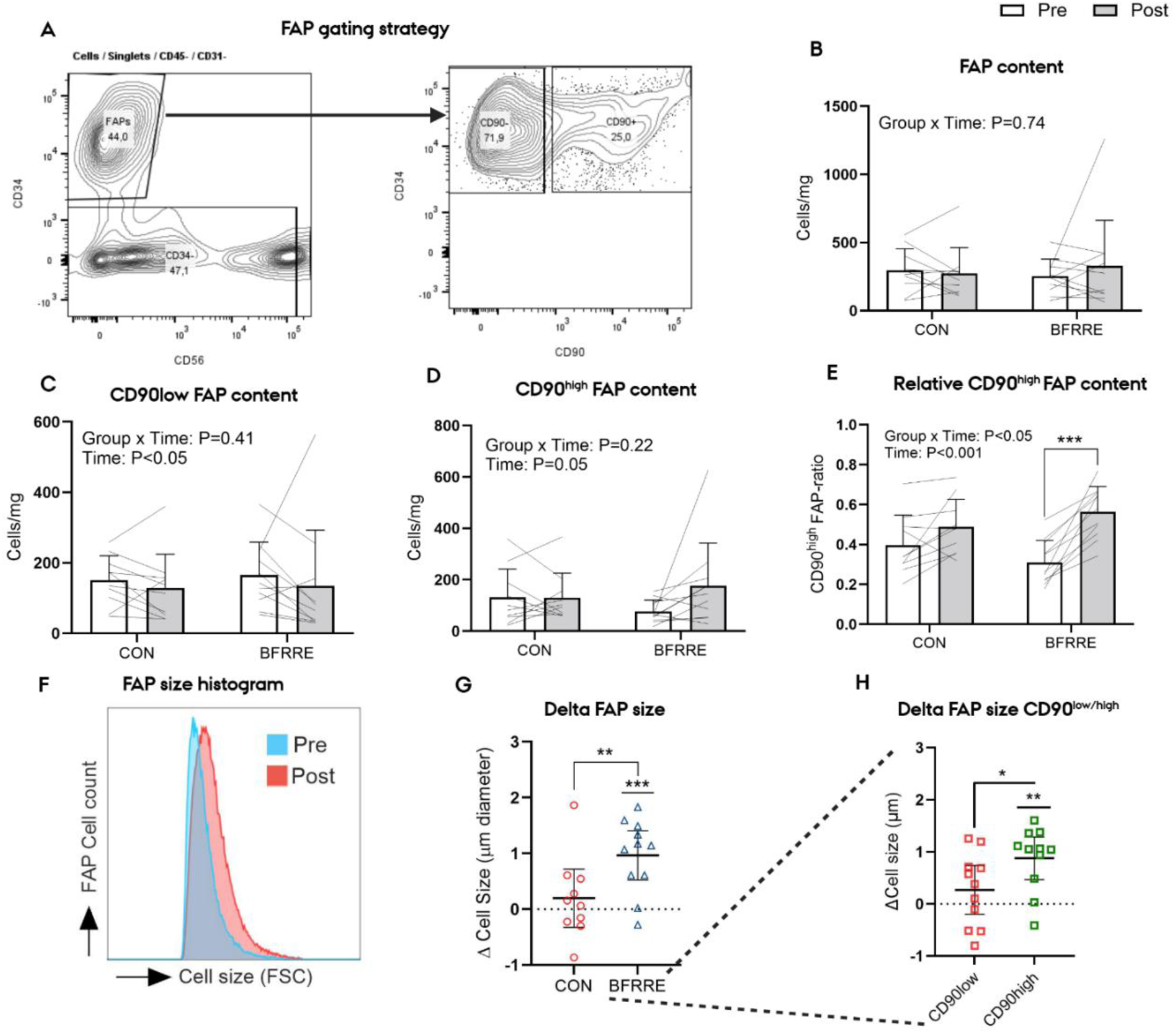
Phenotypic remodeling of FAPs during exercise induced hypertrophy. (A) Representative gating strategy used to identify FAPs (CD45⁻CD31⁻CD34⁺) and subsequently distinguish CD90^low^ and CD90^high^ FAP subpopulations. (B) Total FAP content, (C) CD90^low^ FAP content, and (D) CD90^high^ FAP content before (Pre) and after (Post) the intervention in the CON and BFRRE groups, expressed relative to muscle tissue weight. (E) Relative abundance of CD90^high^ FAPs, expressed as a proportion of the total FAP population. (F) Representative forward-scatter distributions illustrating the change in FAP size from Pre to Post. (G) Pre-to-Post change (Δ) in mean FAP diameter in the CON and BFRRE groups. (H) Pre-to-Post change in the diameter of CD90^low^ and CD90^high^ FAPs within the BFRRE group. Lines in panels B–E connect paired observations from individual participants. Bars and error bars represent mean ± SD; symbols in panels G and H represent individual participants, with horizontal lines and error bars indicating mean ± SD. Group × time interactions and main effects of time are indicated within the panels. *P*<0.05, \**P*<0.01, and \*\**P*<0.001. FAP, fibroadipogenic progenitor; BFRRE, blood flow-restricted resistance exercise; CON, non-exercising control; FSC, forward scatter.

### Gene expression analysis (RNA-sequencing)

The secretome profile of CD90^high^ vs CD90^low^ FAPs was determined by RNA expression analysis. Gene expression data was obtained from a publicly available dataset [32] (EGA accession number: EGAS00001005599). The raw fastq files were processed using Trim Galore (Babraham Bioinformatics) for adapter and quality trimming. Subsequently, gene expression was quantified using Salmon [37]. Predicted secreted genes were acquired from The Protein Atlas [38], and these gene names were used to filter the gene expression data. Differential expression analysis was performed using DESeq2 [39].

### Conditioned media experiments

To investigate the functional implications of FAP subtype composition on skeletal muscle growth, CD90^high^ and CD90^low^ FAP subsets were isolated from skeletal muscle from orthopedic surgeries (n=5), including MuSCs as a cellular control (for donor information see *Supplementary table 2*). Following FACS, 15.000 cells were seeded on ECM-coated 96 well half-area well plates (Corning, USA). Cells were collected and plated in a wash buffer. Twenty-four hours after seeding, the media was aspirated and cells were carefully washed 3x in PBS, before administration of 180 ul wash-buffer. After 48 hours of incubation the media was collected and used for subsequent culture of myotubes. C2C12 myoblasts (#91031101, Sigma-Aldrich, passage 12) were expanded in a T75 culture flask. Upon passaging, the cells were seeded at 5000 cells/well in an ECM-coated 96-well plate. Cells were grown into approx. 70-80% confluency in high glucose (4.5 g/L) DMEM and 20% FBS prior to initiating differentiation by serum starvation (2% HS) in low glucose DMEM. Differentiation media was changed every other day for 10 days. Then, the cells were washed 3 times in PBS, before administration of FAP-conditioned media for 48 hours.

### Immunocytochemistry and imaging

Following fixation in 4% PFA, myotubes permeabilized and blocked with 0.01% triton-X and 10% GS in PBS with 1% BSA. Overnight incubation with MF20 in a 1:10 solution followed by 2 hours of incubation using goat-anti-mouse AF647 secondary antibody was used to visualize myosin heavy chain positive myotubes, which were counterstained with DAPI. Images were acquired of the entire well using the EVOS M700 automated imaging system (Thermo Fisher Scientific). Myotube size was assessed as the average diameter of three randomly selected measurements along the length of each myotube, and the mean myotube diameter in each well was based on normalization to the amount of myotubes quantified. Only myotubes that were entirely visible within each field of view were eligible for analyses. All analysis of myotube size was performed by the same, blinded investigator using ImageJ.

### Macrophage polarization and Multiplex ELISA

To examine the cytokine secretion from polarized macrophages, we isolated monocytes from human PBMCs by the magnetic activated cell sorting (MACS) (Miltenyi Biotech, Sweden). Briefly, 10 mL of blood was collected in EDTA-tubes. Red blood cells were lysed using 1x RBC lysis buffer in sterile water. The suspension was centrifuged at 2000 rpm for 10 minutes, and the supernatant was removed. The pellet was resuspended in 1 mL FACS buffer (1% BSA + 2nM EDTA in PBS) and incubated with 20 µL CD14 microbeads and human Fc Block for 30 minutes at 4 degrees during agitation in accordance with the manufacturers protocol (Milteny Biotech, Sweden). After wash and removal of the supernatant CD14+ monocytes were isolated using the MS-column (130-096-201, Miltenyi Biotech, Sweden). Finally, monocytes were resuspended in macrophage medium (10% FBS + 50 ng/ml M-CSF in 1640 RPMI) and plated at equal densities of 80.000 cells pr. cm^2^ in ECM-coated 48 well plates. Macrophage differentiation was assured by replenishment of the macrophage media using fresh M-CSF every other day for 7 days. Upon differentiation, macrophages were stimulated with either a combination of IL4 and IL13 (both 20 ng/µL) for varying doses of LPS (10, 50 and 100 ng/µL) for anti- and pro-inflammatory differentiation, respectively for 24 hours. Control macrophages remained unstimulated. For the subsequent 24 hours media was conditioned by differentially stimulated macrophages and collected measuring of the concentration of secreted cytokines using multiplex ELISA. The cytokines were assessed using a predefined 17-plex human cytokine assay; IL-1β, IL-2, IL-4, IL-5, IL-6, IL-7, IL-9, IL-10, IL-12p70, IL-13, IL-17A, G-CSF, GM-CSF, MCP-1, MIP-1β, INF-γ and TNF-α (Bio-Rad, Hercules, California, USA), according to the manufacturer’s instructions. The responses were analyzed using the BioPlex Manager 6.0 software (BioRad) and detection limits were between 0.5 and 5 pg/ml.

### Cytokine experiments

The functional implications of cytokine release from pro-inflammatory macrophages were investigated using recombinant human TNF-α on human primary FAPs. FAP proliferation rate was investigated using EdU-incorporation assays analyzed using flow cytometry (Penteon, Novocyte). Here, human primary FAPs from three healthy young donors from a concomitant study [40] were used as model FAPs (for donor characteristics see *Supplementary Table 3*). Following expansion (passaging 3-5 times), cells were seeded at 20.000 cells/cm^2^ in 96 well plates. Twenty-four hours after seeding the cells were treated with a 48-hour EdU-pulse (10uM) co-administrated with recombinant human TNF-α (2-20 ng/mL). EdU and TNF-α were diluted in a mixture of DMEM low and high glucose (ratio 1:1) supplemented with 20% FBS and 1% P/S.

### Statistics

Data were analyzed by a mixed effect linear model with group, time and group x time interaction as factors of interest. When significant group x time interactions were observed, pairwise comparisons were made between groups as well as separate time points (pre vs. post). The model was validated by test for equal standard deviations and visual examination of QQ-plots. Data that did not adhere to a normal distribution was appropriately log-transformed prior to analysis. Normalized gene counts between subpopulation FAPs were compared using paired t-tests. Finally, associations between changes in mononuclear cell content were evaluated using linear regressions. Spearmans correlation coefficient was used to estimate the strength of the association, given the non-gaussian distribution of the absolute changes in cell content. All statistical analyses were conducted using STATA 15.0 (StataCorp, College Station, TX, USA) with an alpha level of 0.05 delineating statistically significant outcomes. Data are graphically presented as individual values and corresponding group means ± SDs. All graphical presentation of data was performed using GraphPad Prism v.7 (GraphPad Software, La Jolla, CA, USA).

## Results

### Alterations in the subcellular composition of FAPs supports exercise-induced skeletal muscle hypertrophy

As previously reported, six weeks BFRRE elicited a 20% increase in mean fiber cross sectional area in the current trial [33], thus providing a context to examine changes in the muscle niche during robust muscle fiber hypertrophy. Using previously established pan-markers (CD34+CD45-CD31-), we did not observe marked increases in overall FAP content with BFRRE compared to control (Fig. 1A). Based on our previous work [32], identifying CD90 as a marker of proliferative and collagen-productive FAPs, we subcategorized FAPs into CD90^low^ and CD90^high^ to investigate subsets of the overall FAP population (Fig. 1B-C). While no group x time interaction was observed for CD90^high^ or CD90^low^ cell content (pr mg of tissue), BFRRE increased the relative abundance of CD90^high^ cells while reducing CD90^low^ cell content (Fig. 1B–C). Normalization of CD90^high^ FAP content to overall FAP content (CD90^high^ FAP-ratio) revealed a marked increase of the proportion of CD90^high^ FAPs selectively in BFRRE group (Fig. 1D), indicating a phenotypical shift in favor of this population. This was associated with a marked increase in FAP cell size (Fig. 1F), that was specifically driven by a selective increase in the cell size of the CD90^high^ FAPs (Fig. 1G). FAP cell size has been linked to increased metabolic and protein synthetic activity, thus the enlargement of CD90^high^ FAPs may reflect a transition towards more actively supporting the requirement for muscle remodeling through protein synthesis and secretion [5, 41]. Given the increased collagen production of CD90^high^ FAPs compared to CD90^low^ this may support remodeling of the ECM in context of marked muscle fiber hypertrophy [42, 43]. To explore potential functional implications of FAP phenotype, we analyzed previously generated RNAseq data from FACS isolated CD90^high^ and CD90^low^ FAPs [32]. A total of 88 genes encoding secreted proteins were differentially expressed between the two populations (*Supplementary Table 1*). Indeed, our transcriptomic data and pathway enrichment analysis revealed CD90^high^ FAPs to be involved collagen organization, formation, and biosynthesis, supporting the contention that a phenotypical shift in favor of CD90^high^ may promote these processes (Supplementary Fig. 2). In addition to their role in collagen production, FAPs are acknowledged as important mediators of cellular crosstalk within the muscle microenvironment through paracrine signaling. Specifically, pre-clinical evidence suggests that FAPs may secrete trophic mediators of muscle fiber growth and degradation, acting through both direct and indirect mechanisms [12, 15, 29, 44]. Based on previous rodent studies [15, 44, 45], we then identified known regulators of muscle mass secreted by FAPs within our data set and directly compared the normalized gene counts between CD90^high^ and CD90^low^ counterparts, including MuSCs as a cellular point of reference (Fig. 2A-G). While no difference was observed for ligands such as IGF1 and follistatin, the expression of GDF10, Wnt-1 inducible secreted protein (WISP1) and thrombospondin (THBS1) were markedly enriched in CD90^high^ compared to CD90^low^ FAPs. The secretion of WISP1 and THBS1 have been identified as central contributors in establishing a transitory niche for mobilization of MuSCs during extensive regeneration and overload-induced hypertrophy though myonuclear donation in mice [15, 29]. Interestingly, GDF10, a member of the TGF-β superfamily, directly stimulates myofiber growth and declining GDF10 levels have been associated with age-related muscle atrophy in humans [45]. Amongst secreted DEGs with a known negative regulatory effect of muscle mass, both tolloid-like protein 2 (TLL2) and inhibin (INHBB) were markedly enriched in CD90^low^ FAPs (Fig. 2F-G, Supplementary table 1). As a secreted metalloprotease, TLL2 is known to activate latent myostatin within the ECM [46]. Similarly, INHBB can bind to activin receptors, hereby promoting SMAD phosphorylation and downstream modulation of genes associated with muscle atrophy [47]. To understand the functional implication of CD90^high^ vs CD90^low^ FAPs in muscle growth, we generated conditioned media from freshly FACS isolated FAPs collected from samples obtained from orthopedic surgeries (Fig. 2 F-G). Here, we found a markedly increased diameter of myotubes upon exposure to CD90^high^ FAP-CM compared to CD90^low^ FAP-CM.

**Fig. 2.**
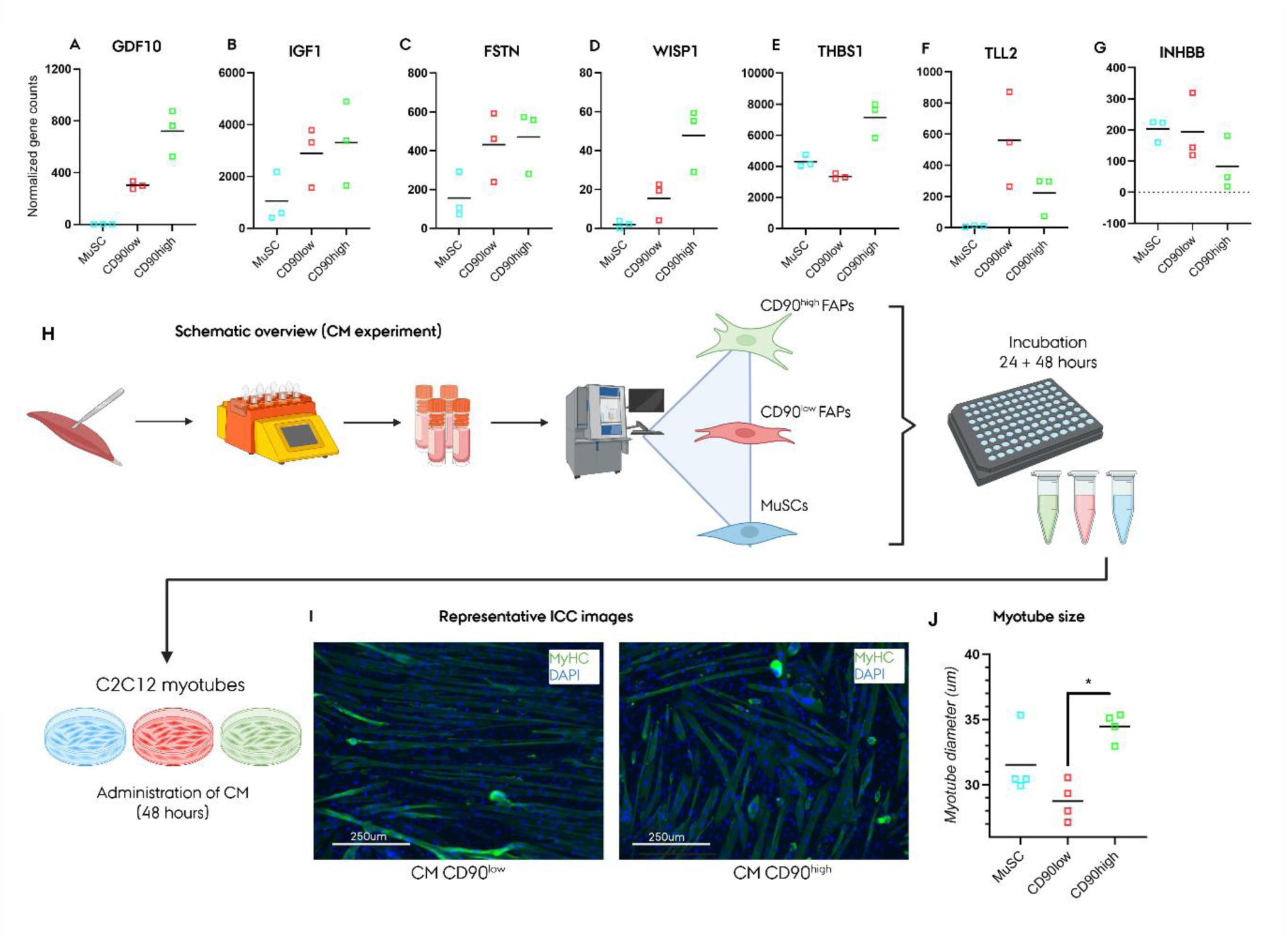
CD90^high^ FAPs exhibit a trophic gene-expression profile and promote myotube growth in vitro. (A–G) Normalized expression of genes encoding growth differentiation factor 10 (GDF10), insulinlike growth factor 1 (IGF1), follistatin (FSTN), WNT1-inducible signaling pathway protein 1 (WISP1), and thrombospondin 1 (THBS1), tolloid-like protein 2 (TLL2), and inhibin (INHBB) in muscle stem cells (MuSCs) and CD90^low^ and CD90^high^ FAPs. Symbols represent individual samples, and horizontal lines indicate group means. (E) Schematic overview of the conditioned-medium (CM) experiment. MuSCs and CD90^low^ and ^CD90high^ FAPs were isolated from human skeletal muscle by fluorescence-activated cell sorting and cultured to generate cell type-specific CM. C2C12 myotubes were subsequently treated with CM for 48 hours. (F) Representative immunocytochemical images of C2C12 myotubes treated with CM from CD90^low^ or CD90^high^ FAPs. Myosin heavy chain (MyHC) is shown in green and nuclei stained with DAPI in blue. Scale bars, 250 µm. (G) Mean diameter of myotubes treated with CM derived from MuSCs, CD90^low^ FAPs, or CD90^high^ FAPs. Symbols represent individual donors/experiments, and horizontal lines indicate group means. *P*<0.05; CM, conditioned medium; DAPI, 4′,6-diamidino-2-phenylindole; FAP, fibroadipogenic progenitor; ICC, immunocytochemistry; MuSC, muscle stem cell.

Collectively, our findings highlight the exercise responsiveness of skeletal muscle CD90^high^ FAPs. Furthermore, the identified functional differences amongst CD90^high^ and CD90^low^ FAPs in providing trophic support for muscle growth, elucidates additional insights into the CD90 subsets and emphasizes the importance of FAPs in governing skeletal muscle adaptations.

### Muscle hypertrophy is associated with and increase in pro-inflammatory macrophages

The skeletal muscle microenvironment is homing various resident and infiltrating immune cells. While transient alterations of immune cell content are clearly important for orchestrating the muscle regenerative response to injury, the regulation of immune cell content in response to prolonged exercise remain sparsely investigated. When examining overall hematopoietic cell content we found no exercise-specific increases in overall CD45+ hematopoietic cells (Fig. 3B). However, we noted a uniform increase in CD45+ cell content across subjects in the BFRRE group. We then focused on the myeloid compartment (CD45+CD14+). Again, while no group x time interaction effects were observed for overall monocyte/macrophage content, the relative individual increases seemed more pronounced and uniform with BFRRE (Fig. 4B). Previously, we confirmed the majority of CD45+CD14+ immune cells in skeletal muscle to co-express classical macrophage markers such as CD68 [35]. To distinguish between inflammatory profiles of macrophages we utilized CD11c as a marker for separating pro- (CD11c+) and anti-inflammatory macrophages (CD11c-). While the increases in both CD11c+ and CD11c-macrophage content seemed to be more pronounced with BFRRE, we found no significant interactions between the groups (Fig. 3C-D). However, the CD11c+/total macrophage-ratio revealed that the proportion of CD11c+ macrophages selectively increased with BFRRE (Fig. 3E). Interestingly, in human skeletal muscle the abundance of pro-inflammatory macrophages seems to decline with age, whereas anti-inflammatory macrophages become more prevalent [48]. Secreted factors from pro- and anti-inflammatory macrophages are known to differentially affect both FAP and MuSC function. For instance, factors secreted from pro-inflammatory macrophages reduces the adipogenic potential of FAPs, whereas anti-inflammatory factors promote it [49]. We speculated if the exercise-induced increase in pro-inflammatory macrophages may support healthy muscle remodeling by limiting the expansion of the FAPs, as observed in preclinical models of muscle injury [10]. To qualify this hypothesis, we generated human macrophages from CD14+ PBMCs in vitro (Fig. 4A). Upon inflammatory polarization of the macrophages we analyzed conditioned media for the concentration of various cytokines (Fig. 4B). Here we found marked increases in the secretion of IL6, IL8, IL10, and TNF-α from pro-inflammatory macrophages, in line with our previously reported pro-inflammatory cytokine profile of CD11c+ macrophages isolated directly from skeletal muscle [35]. Thus, we decided to evaluate the isolated effect of TNF-α on human FAP function in vitro. Using EdU-incorporation, we found that TNF-α markedly reduced the proliferation of FAPs in a dose-depended manner (Fig. 4D). Thus, we speculate that TNF-α mediated signaling from macrophages is an important gatekeeper for preventing excessive FAP accumulation. In support of this mechanistic link, the absolute change in CD11c+ macrophages correlated positively with changes in CD90^high^ FAP content (ρ = 0.69, P < 0.001), but not with changes in CD90^low^ FAP content (ρ = 0.04, P = 0.42), emphasizing the close connection between pro-inflammatory macrophages and FAP subpopulations (Fig. 5).

**Fig. 3.**
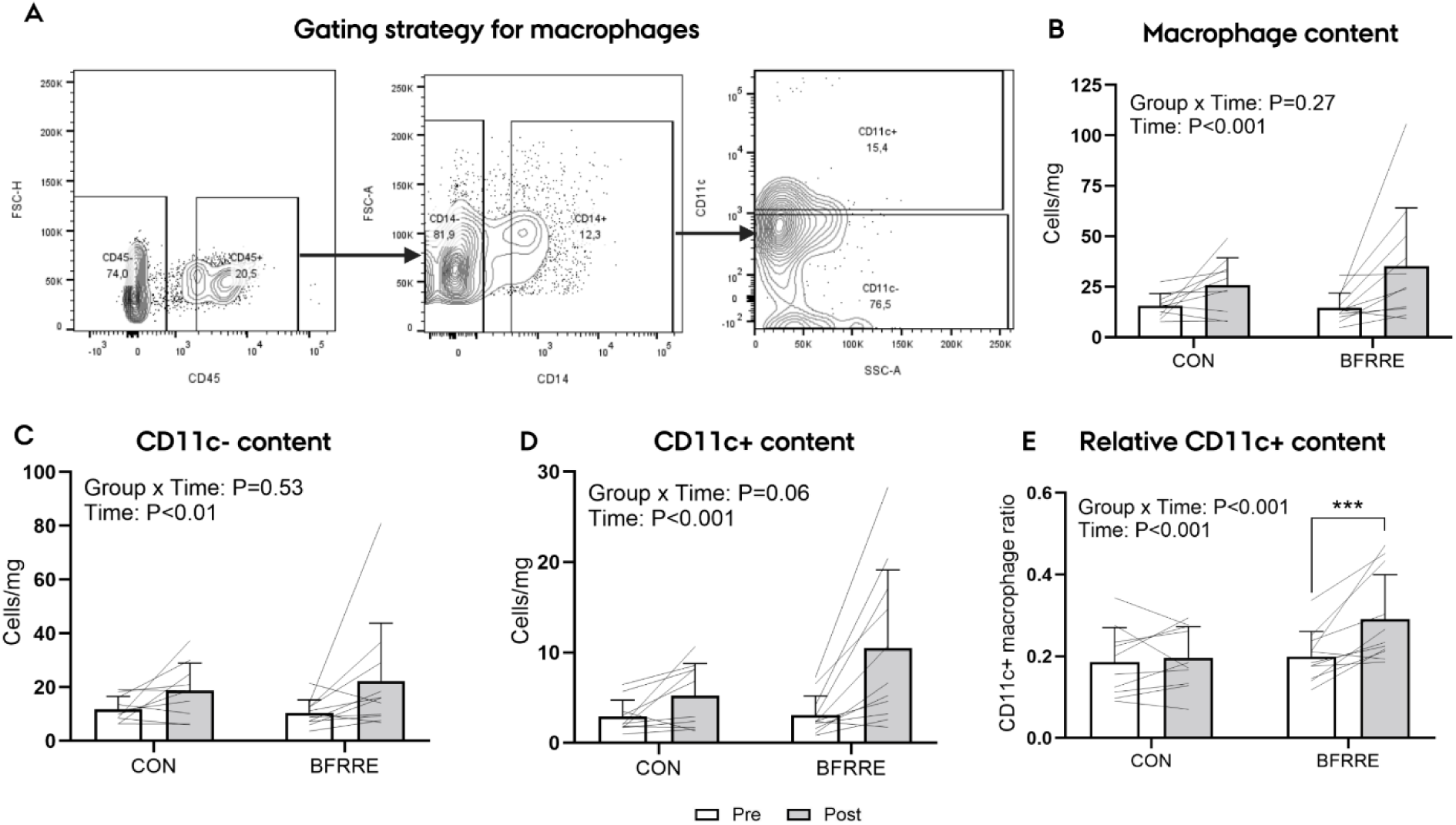
Blood flow-restricted resistance exercise increases the relative abundance of CD11c⁺ skeletal muscle macrophages. A) Representative flow-cytometric gating strategy used to identify CD45⁺CD14⁺ macrophages and subsequently distinguish CD11c⁻ and CD11c⁺ macrophage populations. (B–D) Total macrophage, CD11c⁻ macrophage, and CD11c⁺ macrophage content before (Pre) and after (Post) the intervention in the control (CON) and blood flow-restricted resistance exercise (BFRRE) groups, expressed relative to muscle tissue weight. (E) Relative abundance of CD11c⁺ macrophages, expressed as a proportion of the total macrophage population. Lines connect paired observations from individual participants. Bars and error bars represent mean ± SD. Group × time interactions and main effects of time are indicated within the panels. \*\**P*<0.001. BFRRE, blood flow-restricted resistance exercise; CON, non-exercising control; FSC, forward scatter; SSC, side scatter.

**Fig. 4.**
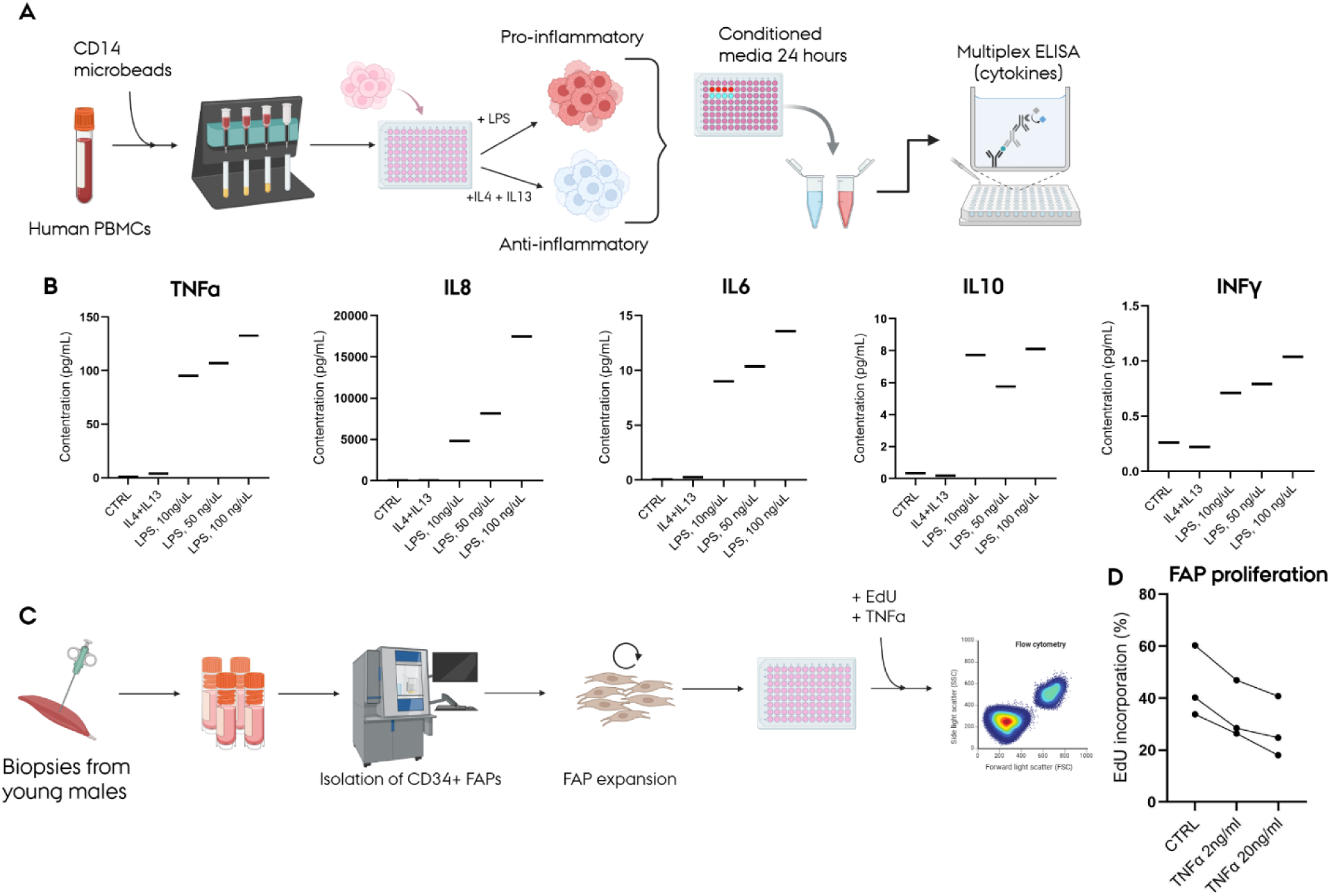
Pro-inflammatory macrophage-derived signals suppress human FAP proliferation. (A) Schematic overview of the macrophage polarization and cytokine-profiling experiment. CD14⁺ monocytes were isolated from human peripheral blood mononuclear cells (PBMCs) using magnetic microbeads and stimulated with lipopolysaccharide (LPS; 10–100 ng/mL) or interleukin IL4 and IL13 to induce pro- or anti-inflammatory phenotypes, respectively. Conditioned media were collected after 24 hours and analyzed using a multiplex cytokine assay. (B) Concentrations of tumor necrosis factor-α (TNF-α), IL8, IL6, IL10, and interferon-γ (IFN-γ) in conditioned media from unstimulated control cells and cells stimulated with IL-4 plus IL-13 or increasing concentrations of LPS. (C) Schematic overview of the FAP proliferation experiment. CD34⁺ FAPs were isolated from skeletal muscle biopsies obtained from young men, expanded in vitro, and treated with TNF-α in the presence of 5-ethynyl-2′-deoxyuridine (EdU). EdU incorporation was subsequently quantified by flow cytometry. (D) Proportion of EdU⁺ FAPs following treatment with vehicle control or TNF-α at 2 or 20 ng/mL. Connected symbols represent cells obtained from the same donor. EdU, 5-ethynyl-2′-deoxyuridine; FAP, fibroadipogenic progenitor; IFN-γ, interferon-γ; IL, interleukin; LPS, lipopolysaccharide; PBMC, peripheral blood mononuclear cell; TNF-α, tumor necrosis factor-α.

**Fig. 5.**
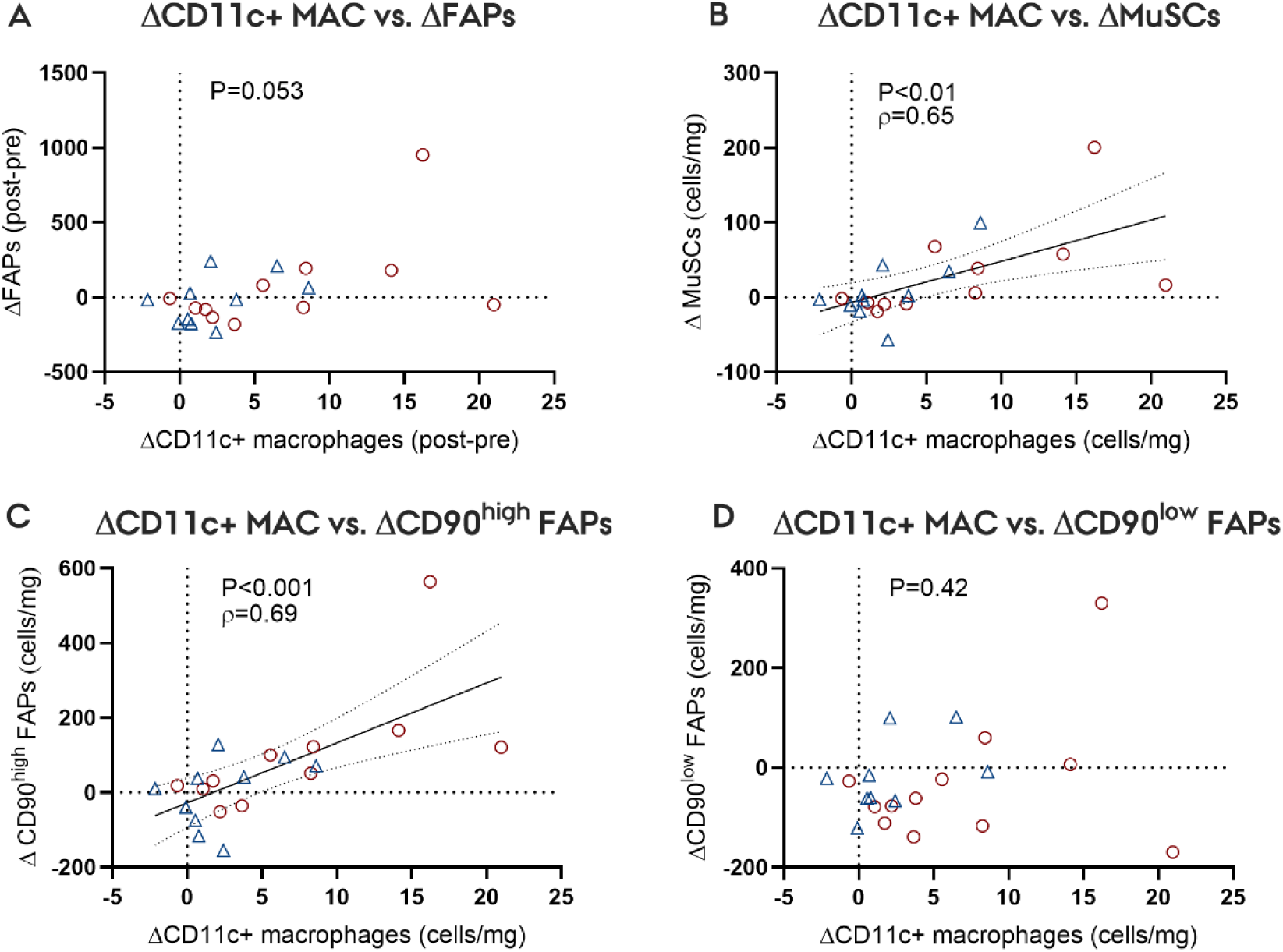
Changes in CD11c⁺ macrophage content are associated with changes in muscle stem cell and CD90^high^ FAP content. Associations between the Pre-to-Post change (Δ) in skeletal muscle CD11c⁺ macrophage content and corresponding changes in (A) total fibroadipogenic progenitor (FAP) content, (B) muscle stem cell (MuSC) content, (C) CD90^high^ FAP content, and (D) CD90^low^ FAP content. Changes are expressed as Post minus Pre and normalized to muscle tissue weight. Blue triangles and red circles represent individual participants in the control (CON) and blood flow-restricted resistance exercise (BFRRE) groups, respectively. Solid lines show fitted regression lines, and dotted lines indicate 95% confidence intervals. Correlation coefficients (Spearmans ρ) and corresponding *P* values are displayed within the panels. BFRRE, blood flow-restricted resistance exercise; CON, non-exercising control; FAP, fibroadipogenic progenitor; MAC, macrophage; MuSC, muscle stem cell.

### Increases in muscle stem cell size accompanies muscle fiber hypertrophy during exercise-induced remodeling

Whereas expansion of the MuSC population is a common observation in response to exercise-induced hypertrophy in humans, we found no apparent increase in MuSC content in the present study. While somewhat unexpected, this was in line with our prior results from immunohistochemistry analysis [33]. However, as was the case for FAPs, we did note a marked increase in MuSC size with BFRRE compared to baseline (Fig. 6 B-C). While no group x time interaction was observed for increases in MuSC size, the selective increase within the BFRRE group does suggest exercise to alter the metabolic status of the cells. Interestingly, our findings of exercise-induced increases in MuSC size mirrors evidence from mice, indicating that prolonged exercise restores age-related impairments in MuSC function, rather than increasing content during skeletal muscle adaptation [25].

**Fig. 6.**
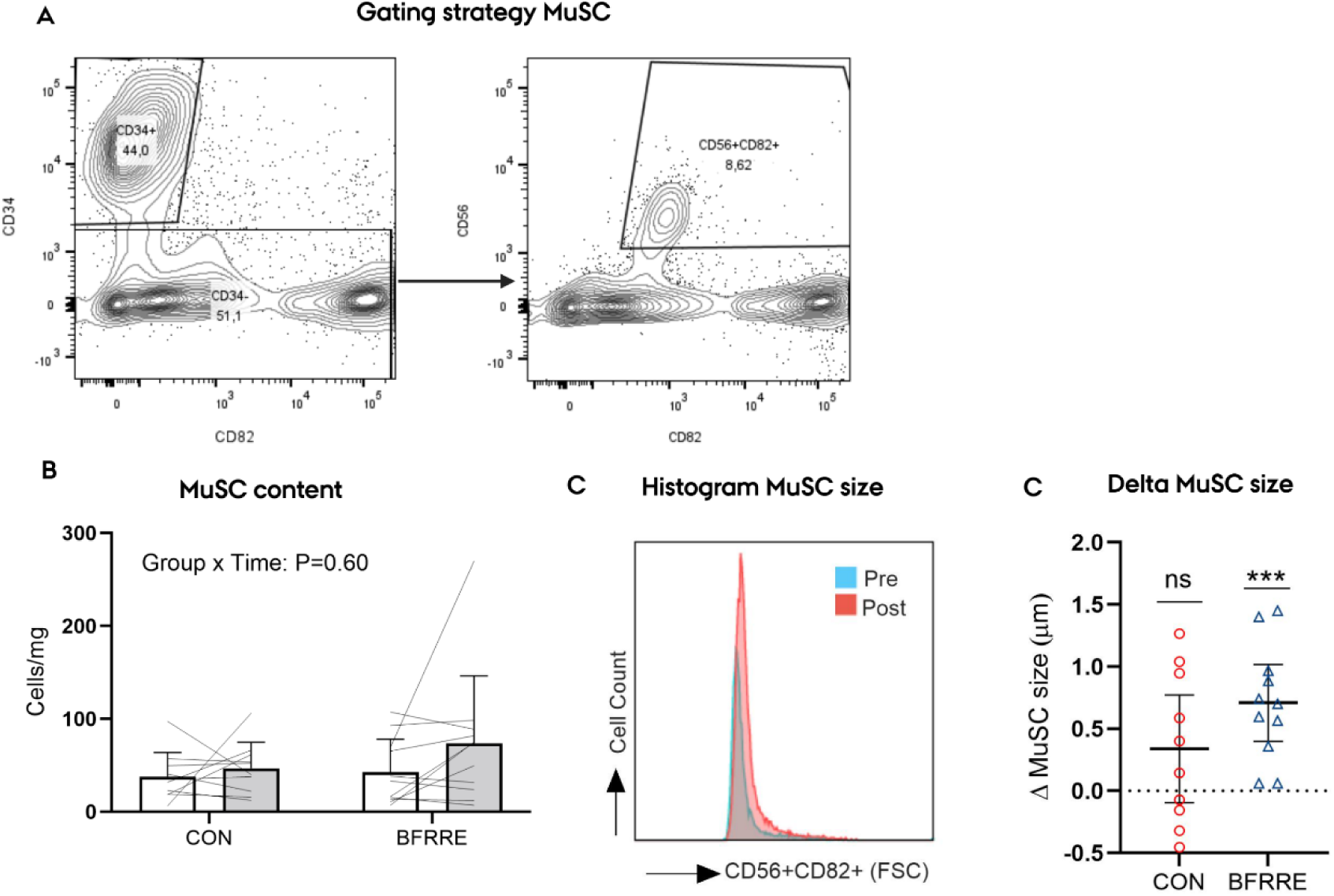
Blood flow-restricted resistance exercise increases skeletal muscle stem cell size without altering muscle stem cell content. (A) Representative flow-cytometric gating strategy used to identify muscle stem cells (MuSCs) as CD45⁻CD31⁻CD34⁻CD56⁺CD82⁺ cells. (B) MuSC content before (Pre) and after (Post) the intervention in the control (CON) and blood flow-restricted resistance exercise (BFRRE) groups, expressed relative to muscle tissue weight. Lines connect paired observations from individual participants. Bars and error bars represent mean ± SD. The group × time interaction is indicated within the panel. (C) Representative forward-scatter distributions illustrating the change in MuSC size from Pre to Post. (D) Pre-to-Post change (Δ) in mean MuSC diameter in the CON and BFRRE groups. Symbols represent individual participants, and horizontal lines and error bars indicate mean ± SD. \*\**P*<0.001; significant within group compared to baseline. BFRRE, blood flow-restricted resistance exercise; CON, non-exercising control; FSC, forward scatter; MuSC, muscle stem cell.

## Discussion

Opposed to our initial hypothesis, we did not observe an increase in overall FAP content following exercise-induced hypertrophy in humans. Instead, we found changes in the phenotypical composition of the FAP pool, which may contribute to exercise-induced remodeling. Specifically, the proportion of CD90^high^ FAPs increased markedly in the context of muscle fiber hypertrophy. This was associated with an overall increase in FAP cell size that was primarily attributed to enlargement of the CD90^high^ population specifically. Our conditioned media experiments further demonstrated functional differences between the FAP subpopulations, as CD90^high^ FAPs provided greater trophic support for myotube growth *in vitro*, compared to CD90^low^ counterparts. Finally, during muscle hypertrophy, the proportion of pro-inflammatory CD11c+ macrophages increased within the muscle microenvironment, which may serve as a counterregulatory mechanism to prevent excessive CD90^high^ FAP expansion, indicated by a direct inhibitory effect of TNF-α on FAP proliferation ex vivo.

Whereas CD90 was initially used as a putative FAP marker, it is now known to characterize a highly proliferative and collagen-productive subset of FAPs [32]. Given the implication of CD90^high^ FAP accumulation in pathological degeneration of skeletal muscle in type II diabetes [32], the exercise-induced increase in CD90^high^/FAP-ratio in the present study may seem counterintuitive. However, while uncontrolled expansion of FAPs (and perhaps particularly CD90^high^ subsets) can certainly drive fibrosis, more controlled expansion may be favorable in context of muscle growth and the required ECM remodeling. Thus, the increased CD90^high^/FAP-ratio may reflect an increased need for collagen production in support of the ECM remodeling associated with muscle fiber hypertrophy [43]. Our transcriptomic data supports this contention, with secreted DEGs related to ECM remodeling, including COL1A1, COL4A1 and FN1. Furthermore, periostin (POSTN), which is associated with activated matrix-remodeling FAP subsets [50, 51] was markedly enriched in CD90^high^ FAPs. Finally, our pathway enrichment analysis consistently identified pathways suggesting that CD90^high^ FAPs may contribute to ECM remodeling (see supplementary Fig. 2).

Interestingly, our findings are corroborated by other studies reporting marked increases in the number of proliferating CD90^+^ interstitial cells in association with exercise-induced hypertrophy in humans [52]. Recently, the FAP content (i.e. PDGFRa+) was shown to be unaffected by prolonged resistance exercise in untrained college-aged males [53]. Collectively, these findings highlight the functional implications of phenotypically distinctive subsets of FAPs and suggests the CD90^high^ subpopulation as a potential meditator of exercise-induced adaptations in humans. In keeping with this, the overall increase in FAP size following BFRRE was largely driven by cellular enlargements preferential to the CD90^high^ population. Previously, overload-induced hypertrophy in mice has been associated with concomitant increases in FAP content and size. Furthermore, it was shown that compensatory fiber hypertrophy was abolished upon FAP depletion, which was partly ascribed to a lack of FAP-secreted THBS1 [29]. Although speculative, the exercise-induced shift toward CD90^high^ FAPs could potentially contribute to this response, consistent with the reported differences in THBS1 expression between FAP subtypes.

Age-dependent declines in GDF10 levels have been shown to impair both muscle anabolic signaling and muscle regeneration in rodent models [15, 45, 54], and have been directly associated with the development of sarcopenia in humans [45]. Notably, among skeletal muscle-resident cell types, the expression of GDF10 is specific to FAPs [55]. Our transcriptomic analysis identified GDF10 amongst the differentially expressed secreted factors from CD90^high^ versus CD90^low^ FAPs. Conversely, the transcription of pro-catabolic ligands associated with activation of latent myostatin and activin/SMAD3-signalling (i.e. TLL2 and INHBB) were both enriched in CD90^low^ FAPs. Our conditioned-media experiment cannot distinguish whether the observed effect on myotube size was driven by pro-anabolic factors by CD90^high^ FAPs or pro-catabolic factors released from CD90^low^ FAPs.

However, it clearly demonstrates that CD90^high^ FAPs provide a more growth-supportive trophic environment for myotubes compared to CD90^low^. Interestingly, these observations are consistent with our recent observations that muscle wasting in cancer cachexia is associated with marked reductions in CD90^high^ FAPs, which were associated with reduced trophic pro-myogenic support in vitro [17]. In this context, we speculate that an increase in the CD90^high^ FAP population may represent one component of exercise adaptation in the aging cellular microenvironment to support exercise-induced remodeling. This speculation warrants future cross-sectional studies directly comparing the composition of subpopulation FAPs across different age groups, to ascertain exercise-associated changes reflect reversal of age-related deficits.

We found that the proportion of CD11c+ pro-inflammatory macrophages substantially increased during muscle hypertrophy. It remains unclear whether this altered macrophage composition is related to polarization of tissue-resident macrophages or increased infiltration of monocytes. Similarly, we cannot exclude that BFRRE itself may have elicited transient inflammatory responses. However, given the duration of the intervention (i.e. six weeks) and the relatively low mechanical stress of the exercise stimulus (i.e. low relative intensity), substantial muscle damage resulting from the final exercise session is unlikely to have occurred [56, 57]. Interestingly, skeletal muscle aging is associated with reductions in overall macrophage content primarily caused by a preferential loss of pro-inflammatory macrophages [48]. Furthermore, an inability to recruit pro-inflammatory macrophages has been proposed to partly explain the attenuation of muscle regeneration and adaptation with increasing age [58]. Thus, the marked increase in the proportion of CD11c+ macrophages observed during muscle growth, may indicate an exercise-associated shift toward a more pro-inflammatory macrophage profile and support a contention that can counter age-related inflammatory imbalance of cellular microenvironment. The exact mechanism driving this and the role of these pro-inflammatory macrophages under homeostatic conditions currently remains unclear. However, pro-inflammatory cytokine production, and particularly TNF-α, has been shown to prevent excessive fibrosis by harnessing FAP expansion during muscle regeneration [10]. Whereas Lemos et al., suggested TNF-α mediated apoptosis to prevent excessive FAP expansion, we found that TNF-α markedly reduced human FAP proliferation. Given the apparent increase in CD90^high^ FAP proportion observed following BFRRE, we speculate that the increase in CD11c+ macrophages serves to prevent unintended accumulation of FAPs during the remodeling process. As such, maintaining inflammatory balance of the skeletal muscle microenvironment may explain the antifibrotic effects of regular exercise [26].

In agreement with our previous reporting of Pax7+ MuSCs on muscle cross sections from the present study [33], BFRRE did not alter the MuSC content when analyzed using flow cytometry. However, BFRRE was associated with a marked increase in the flow-mediated cell size of MuSCs. Indeed, using electron microscopy, increases in MuSC size have previously been reported following prolonged resistance exercise in older individuals [59]. Similarly, voluntary wheel running in old mice was shown to markedly increase MuSC size resulting in an augmented activation capacity *ex vivo*, hereby mitigating age-related deficits in rate of activation [25]. Similarly, we speculate that the increased MuSC size could resemble that of alerted MuSCs primed for activation [60]. Although speculative, MuSC size is closely linked to rate of activation and previous observations that resistance exercise training augments MuSC recruitment in response to single-bout exercise provide indirect support for this speculation [61, 62].

Collectively, our findings shed new light on the coordinated regulation of the cellular microenvironment during skeletal muscle hypertrophy in humans. We highlight dynamic alterations within the heterogenous pool of resident FAPs and macrophages during tissue remodeling in response to exercise, together with changes in MuSC size. These combined cellular adaptations may contribute to a more growth-supportive microenvironment of the muscle and represent mechanisms through which exercise can counteract age-associated features of muscle dysfunction.

### Limitations

Several considerations define the scope of the present findings. The robust myofiber hypertrophy induced by BFRRE provided a well-controlled and biologically relevant model in which to investigate remodeling of the human skeletal muscle microenvironment. However, BFRRE combines mechanical loading, fatigue, ischemia, and metabolite accumulation in a manner distinct from conventional resistance exercise [63]. Thus, the findings reported here should not be assumed to generalize to other exercise modalities without direct investigation. Our phenotypic and morphological analyses did not include paired assessments of intrinsic FAP and MuSC function before and after training. Functional measures—including proliferation, differentiation, and lineage propensity—will therefore be important for determining whether the observed cellular changes translate into functional capacity in vivo. Importantly, however, the conditioned-media experiments provide direct functional evidence that CD90^high^ and CD90^low^ FAPs exert distinct effects on myotube growth. Although the factors responsible for this difference remain to be verified, these experiments extend the observational findings by demonstrating subtype-specific trophic activity ex vivo.

## Acknowledgement

The authors thank Lotte Ahrentoft, Magnus Brandbyge, Gitte Kaiser Hartvigsen, Janni Mosgaard Jensen for their technical and scientific contributions to the work. Florescence activated cell sorting was performed at the FACS Core unit at Aarhus University, Department of Biomedicine, Aarhus, Denmark, and they are likewise thanked for their expertise.

## Supplementary Figs

**Supplementary Fig. 1.**
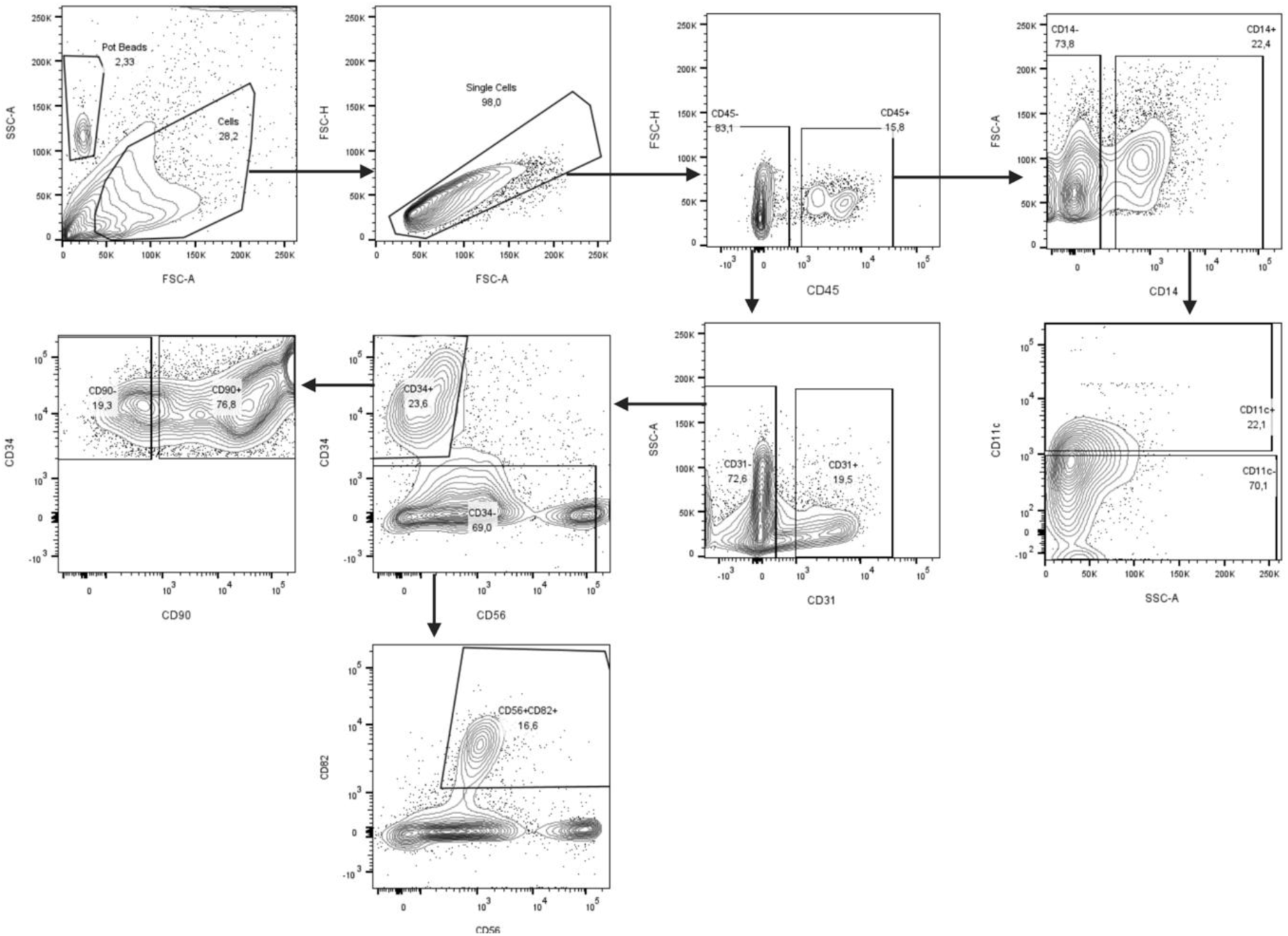
Generalized gating strategy for flow analysis.

**Supplementary Fig. 2.**
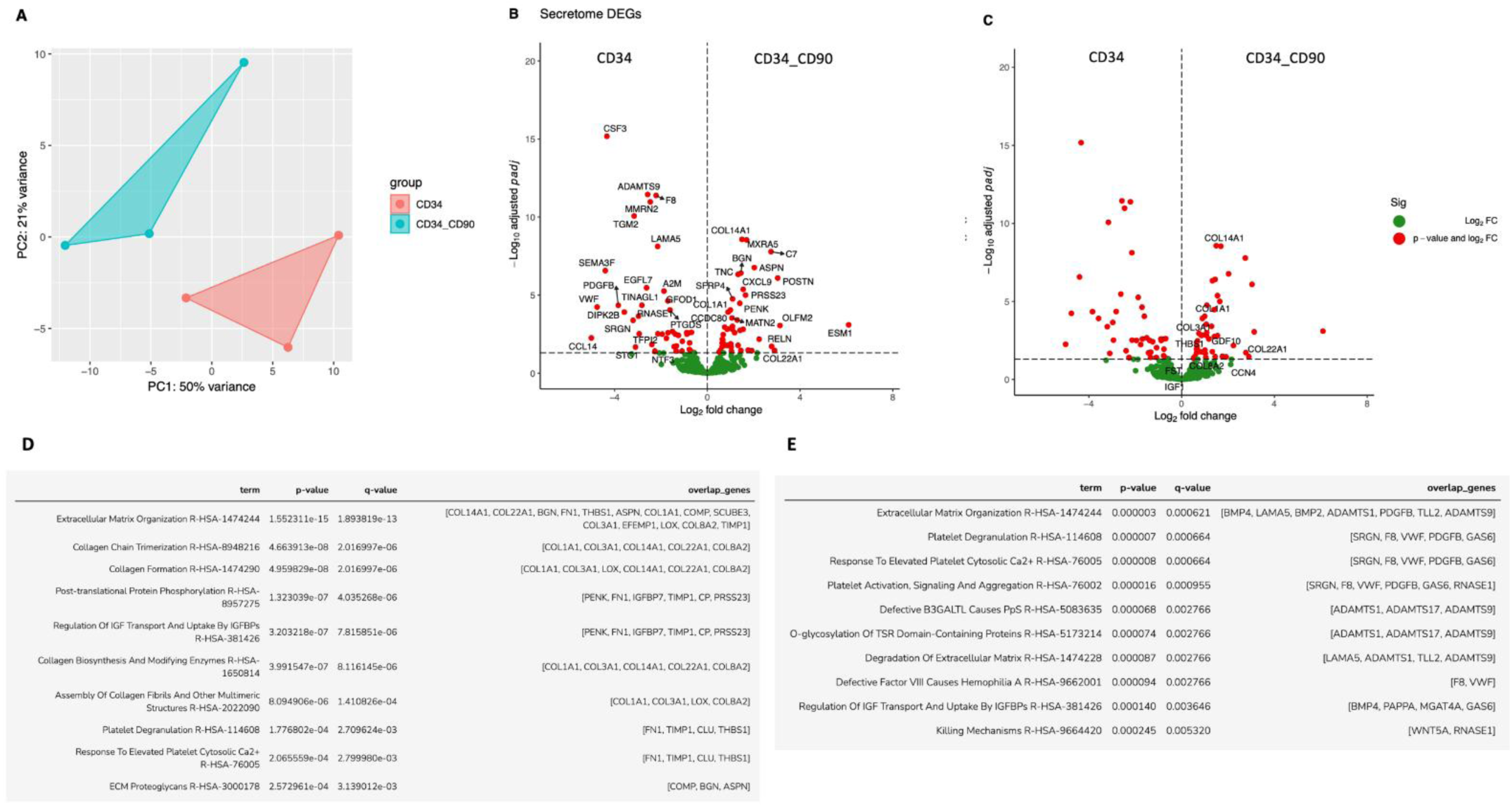
A) Principal Component Analysis (PCA) demonstrated distinct clustering of the two cell types based on the expression of secreted genes. B) Differential expression analysis revealed 88 genes with significant differential expression. C) Changes in selected genes: GDF10, COL22A1, COL14A1, COL8A2, COL1A1, COL3A1, IGF1, FST, CCN4 (Corresponding to WISP1), THBS1. D) Pathway enrichment analysis (REACTOME) was performed on the upregulated secretome genes in CD90^low^ vs CD90^high^ FAPs. E) Pathway enrichment analysis (REACTOME) was performed on the upregulated secretome genes in CD90^high^ vs CD90^low^ FAPs.

**Supplementary table 1.** List of genes (DEGs) in the secretome of CD90^high^ FAPs versus CD90^low^ FAPs. Positive log2(fold)change indicates that genes are upregulated in CD90^high^ FAPs. Adjusted P-values for the comparison is given in the second column. Known positive regulators of muscle mass are highlighted in yellow.

secreted\_DEGs
| log2FoldChange | padj | SYMBOL |
| --- | --- | --- |
| 6.094396167579340 | 8.07596079401837E-04 | ESM1 |
| 3.1312530884635600 | 8.94287388265173E-04 | OLFM2 |
| 3.0354268648449600 | 8.0835713663165E-07 | POSTN |
| 2.9002565316409300 | 0.03495119853810560 | COL22A1 |
| 2.7702779630236100 | 0.019221475848246800 | RELN |
| 2.7462998464110200 | 1.65943954110219E-08 | C7 |
| 2.235009732975950 | 0.006675775541446390 | SCUBE3 |
| 2.0262483503083900 | 1.72613732689164E-07 | ASPN |
| 1.9182346501411700 | 0.03631136804040350 | C16orf89 |
| 1.758010411017330 | 0.03341440766759960 | C1QTNF4 |
| 1.6920545320118000 | 2.90581288292958E-09 | MXRA5 |
| 1.6461566890660100 | 1.00469170816758E-05 | PRSS23 |
| 1.5458153433504800 | 4.30218470767188E-06 | CXCL9 |
| 1.5455342954454200 | 0.0015681531421857300 | COMP |
| 1.4917882343716500 | 2.71040265090942E-09 | COL14A1 |
| 1.4456376181937200 | 3.89804935087686E-07 | BGN |
| 1.4448076161777600 | 0.03495119853810560 | CTHRC1 |
| 1.4113225771507700 | 0.0018046738812791000 | CP |
| 1.4000631917115100 | 3.36408634300374E-05 | PENK |
| 1.3278551948689900 | 4.73051075355825E-07 | TNC |
| 1.3232209578252300 | 0.016642791499449100 | IGSF10 |
| 1.2874701716397900 | 3.94082533639952E-04 | CLU |
| 1.1758993743580100 | 0.0025631462098481200 | FAM180B |
| 1.089861101705930 | 1.7532811003621E-05 | SFRP4 |
| 1.0838802975928900 | 0.0010095121959790100 | SFRP2 |
| 1.0756631425940200 | 0.0015485975121785600 | GDF10 |
| 1.0619199096133900 | 2.94218260441986E-04 | MATN2 |
| 1.041147593999270 | 0.03953088908462930 | COL8A2 |
| 1.0194676944119800 | 0.021354538205065500 | ADAMTSL1 |
| 1.0074914052444200 | 0.013296107847785900 | OGN |
| 0.9947482539799510 | 9.02410812133004E-05 | COL1A1 |
| 0.8944468801328790 | 1.2342304099419E-04 | CCDC80 |
| 0.8825898583789050 | 0.0014635868453425600 | COL3A1 |
| 0.8567573638382610 | 0.0170482691785997 | LOX |
| 0.8555943123657040 | 0.0015406651540623100 | SOD3 |
| 0.7450492205560470 | 0.0011304498115762900 | FN1 |
| 0.7278155679630060 | 0.016642791499449100 | THBS1 |
| 0.68334337293418 | 0.0034145889110666200 | AEBP1 |
| 0.6756478395308220 | 0.011503033099509600 | MGP |
| 0.6670071852966750 | 0.007453840490933120 | IGFBP7 |
| 0.647453451718022 | 0.006675775541446390 | HMCN1 |
| 0.6379909596597490 | 0.020016589637998100 | TIMP1 |
| 0.6373578642991610 | 0.011503033099509600 | CFH |
| 0.5703532108398920 | 0.03341440766759960 | EFEMP1 |
| 0.514496039101532 | 0.043350422442536200 | ABI3BP |
| -0.5669053636157230 | 0.048223349470347100 | WNT5A |
| -0.6875646168206250 | 0.0024379011964912200 | HTRA3 |
| -0.7143528607040760 | 0.0386466834991489 | BMP2 |
| -0.7634722737863940 | 0.011593033823459300 | ADAMTS1 |
| -0.7755159336252630 | 0.0305775632367937 | GAS6 |
| -0.8079199876521770 | 0.0026530071087225200 | BMP4 |
| -0.8821059945222890 | 0.003172712959492760 | HMCN2 |
| -0.8922265440091740 | 0.0025631462098481200 | BMPER |
| -1.0782390136086300 | 0.009186159410308070 | PAPPA |
| -1.0914804608488000 | 0.0386466834991489 | CRLF1 |
| -1.2472153790053300 | 0.0038257410477062000 | BMP6 |
| -1.3314627916451400 | 0.017973429355028700 | CBLN4 |
| -1.342346770537560 | 0.0386466834991489 | DKK2 |
| -1.3825663555469100 | 0.0031589668334896000 | ADAMTS17 |
| -1.4338182081178800 | 0.020016589637998100 | EFNA1 |
| -1.5052105275434200 | 0.0021556339161538000 | OLFM4 |
| -1.610752007377230 | 9.02410812133004E-05 | PTGDS |
| -1.6959315863645600 | 0.0025631462098481200 | TLL2 |
| -1.707393314239140 | 2.36376752712853E-05 | GFOD1 |
| -1.7666137838908100 | 0.0058608030007520100 | INHBB |
| -1.8732088251679200 | 5.55542416152889E-06 | A2M |
| -1.9497960417845000 | 0.0031589668334896000 | SEMA3G |
| -2.1368577988260700 | 0.0030352173703510600 | LPL |
| -2.145386591681830 | 7.62752469427348E-09 | LAMA5 |
| -2.2097616762039100 | 4.15916193918866E-12 | F8 |
| -2.2656000384052700 | 0.0386466834991489 | NTF3 |
| -2.2657281739443200 | 0.04152372376582560 | VSTM2A |
| -2.3947343916195500 | 0.014624407563684300 | MGAT4A |
| -2.4612984056335400 | 1.0669236899422E-11 | MMRN2 |
| -2.5748954564146800 | 3.59108677632325E-12 | ADAMTS9 |
| -2.6239905749777500 | 3.39450936713917E-06 | EGFL7 |
| -2.824378077920100 | 4.48137257783827E-05 | TINAGL1 |
| -2.9411815074975000 | 0.0030352173703510600 | TFPI2 |
| -2.9698254473555100 | 2.23001083446102E-04 | RNASE1 |
| -3.0995596706919600 | 0.021403917233691700 | STC1 |
| -3.1598784417287400 | 8.59521238791364E-11 | TGM2 |
| -3.2054204656351300 | 4.1505776692023E-04 | SRGN |
| -3.5827862264756100 | 1.2342304099419E-04 | DIPK2B |
| -3.8428722655497100 | 4.54385926735081E-05 | PDGFB |
| -4.33458240449627 | 6.73204625245189E-16 | CSF3 |
| -4.4034606726034800 | 2.71034609264272E-07 | SEMA3F |
| -4.754586098334750 | 5.82620054274377E-05 | VWF |
| -5.003475072298860 | 0.005618393771806230 | CCL14 |

**Supplementary table 2.** Tabel depicting the sex, age and clinical indication for orthopedic surgery. Samples collected during surgeries were used generate model FAPs for conditioned media experiments.

| Sex | Age | Clinical indication |
| --- | --- | --- |
| Male | 79 | Ischemia |
| Male | 68 | Ischemia |
| Male | 72 | Ischemia |
| Male | 75 | Ischemia |

**Supplementary table 3.** Tabel depicting the gender, age, height, weight and BMI of subjects that provided biobanked model FAPs for EdU-experiments to assess the effect of TNFα.

| Sex | Age | Height (cm) | Weight (kg) | BMI (kg/m<sup>2</sup>) |
| --- | --- | --- | --- | --- |
| Male | 23 | 189.9 | 81.2 | 22.52 |
| Male | 25 | 192.5 | 88.3 | 23.83 |
| Male | 24 | 188.8 | 78.7 | 22.07 |

## References

1. Visser, M., et al., Muscle mass, muscle strength, and muscle fat infiltration as predictors of incident mobility limitations in well-functioning older persons. J Gerontol A Biol Sci Med Sci, 2005. 60(3): p. 324–33.

2. Vainshtein, A., et al., The hallmarks of skeletal muscle health. Nature Metabolism, 2026.

3. Cruz-Jentoft, A.J., et al., Sarcopenia: revised European consensus on definition and diagnosis. Age Ageing, 2019. 48(4): p. 601.

4. Chakkalakal, J.V., et al., The aged niche disrupts muscle stem cell quiescence. Nature, 2012. 490(7420): p. 355–60.

5. Liu, X., et al., Mesenchymal Stromal Cell-Mediated Intercellular Communication: Mapping the Interactome for Skeletal Muscle Homeostasis and Regeneration. Advanced Science, 2026.

6. Fukada, S.I. and A. Uezumi, Roles and heterogeneity of mesenchymal progenitors in muscle homeostasis, hypertrophy, and disease. Stem Cells, 2023.

7. Uezumi, A., et al., Fibrosis and adipogenesis originate from a common mesenchymal progenitor in skeletal muscle. J Cell Sci, 2011. 124(Pt 21): p. 3654–64.

8. Joe, A.W., et al., Muscle injury activates resident fibro/adipogenic progenitors that facilitate myogenesis. Nat Cell Biol, 2010. 12(2): p. 153–63.

9. Wosczyna, M.N. and T.A. Rando, A Muscle Stem Cell Support Group: Coordinated Cellular Responses in Muscle Regeneration. Dev Cell, 2018. 46(2): p. 135–143.

10. Lemos, D.R., et al., Nilotinib reduces muscle fibrosis in chronic muscle injury by promoting TNF-mediated apoptosis of fibro/adipogenic progenitors. Nat Med, 2015. 21(7): p. 786–94.

11. Chazaud, B., Inflammation and Skeletal Muscle Regeneration: Leave It to the Macrophages! Trends Immunol, 2020. 41(6): p. 481–492.

12. Lukjanenko, L., et al., Loss of fibronectin from the aged stem cell niche affects the regenerative capacity of skeletal muscle in mice. Nature Medicine, 2016. 22(8): p. 897–905.

13. Schüler, S.C., et al., Extensive remodeling of the extracellular matrix during aging contributes to age-dependent impairments of muscle stem cell functionality. Cell Rep, 2021. 35(10): p. 109223.

14. Brorson, J., et al., Complementing muscle regeneration-fibro-adipogenic progenitor and macrophage-mediated repair of elderly human skeletal muscle. Nat Commun, 2025. 16(1): p. 5233.

15. Lukjanenko, L., et al., Aging Disrupts Muscle Stem Cell Function by Impairing Matricellular WISP1 Secretion from Fibro-Adipogenic Progenitors. Cell Stem Cell, 2019. 24(3): p. 433–446 e7.

16. Kajabadi, N., et al., Activation of beta-catenin in mesenchymal progenitors leads to muscle mass loss. Dev Cell, 2023. 58(6): p. 489–505 e7.

17. Battey, E., et al., Pathophysiological remodeling of the skeletal muscle microenvironment in patients with lung cancer. bioRxiv, 2025: p. 2025.11.19.689342.

18. Narasimhan, A., et al., Cachexia-induced alterations of miR-27a-3p drive cell-type specific effects in FAPs and tumor cells that coincide with muscle wasting. Cell Rep, 2026. 45(6): p. 117362.

19. D’Lugos, A.C., et al., Complement pathway activation mediates pancreatic cancer–induced muscle wasting and pathological remodeling. Journal of Clinical Investigation, 2025. 135(12).

20. Wosczyna, M.N., et al., Mesenchymal Stromal Cells Are Required for Regeneration and Homeostatic Maintenance of Skeletal Muscle. Cell Rep, 2019. 27(7): p. 2029–2035 e5.

21. Roberts, E.W., et al., Depletion of stromal cells expressing fibroblast activation protein-alpha from skeletal muscle and bone marrow results in cachexia and anemia. J Exp Med, 2013. 210(6): p. 1137–51.

22. Natarajan, A., D.R. Lemos, and F.M. Rossi, Fibro/adipogenic progenitors: a double-edged sword in skeletal muscle regeneration. Cell Cycle, 2010. 9(11): p. 2045–6.

23. Csapo, R., M. Gumpenberger, and B. Wessner, Skeletal Muscle Extracellular Matrix - What Do We Know About Its Composition, Regulation, and Physiological Roles? A Narrative Review. Front Physiol, 2020. 11: p. 253.

24. Egan, B. and J.R. Zierath, Exercise metabolism and the molecular regulation of skeletal muscle adaptation. Cell Metab, 2013. 17(2): p. 162–84.

25. Brett, J.O., et al., Exercise rejuvenates quiescent skeletal muscle stem cells in old mice through restoration of Cyclin D1. Nat Metab, 2020. 2(4): p. 307–317.

26. Tuñón-Suárez, M., et al., Exercise Training to Decrease Ectopic Intermuscular Adipose Tissue in Individuals With Chronic Diseases: A Systematic Review and Meta-Analysis. Physical Therapy, 2021. 101(10).

27. Wei, W., et al., Organism-wide, cell-type-specific secretome mapping of exercise training in mice. Cell Metab, 2023. 35(7): p. 1261–1279.e11.

28. Kang, X., et al., Exercise-induced Musclin determines the fate of fibro-adipogenic progenitors to control muscle homeostasis. Cell Stem Cell, 2024. 31(2): p. 212–226.e7.

29. Kaneshige, A., et al., Relayed signaling between mesenchymal progenitors and muscle stem cells ensures adaptive stem cell response to increased mechanical load. Cell Stem Cell, 2021.

30. Langston, P.K., et al., Piezo1-dependent activation of stromal cells ignites muscle inflammation in exercise and injury and is associated with inflammaging. Nat Immunol, 2026. 27(3): p. 543–555.

31. Saito, Y., et al., Exercise enhances skeletal muscle regeneration by promoting senescence in fibro-adipogenic progenitors. Nature Communications, 2020. 11(1).

32. Farup, J., et al., Human skeletal muscle CD90(+) fibro-adipogenic progenitors are associated with muscle degeneration in type 2 diabetic patients. Cell Metab, 2021. 33(11): p. 2201–2214 e11.

33. Wang, J., et al., Low-load blood flow-restricted resistance exercise produces fiber type-independent hypertrophy and improves muscle functional capacity in older individuals. J Appl Physiol (1985), 2023. 134(4): p. 1047–1062.

34. Bergstrom, J., Percutaneous needle biopsy of skeletal muscle in physiological and clinical research. Scand J Clin Lab Invest, 1975. 35(7): p. 609–16.

35. Jensen, J.B., et al., Isolation and characterization of muscle stem cells, fibro-adipogenic progenitors, and macrophages from human skeletal muscle biopsies. Am J Physiol Cell Physiol, 2021. 321(2): p. C257–C268.

36. Billeskov, T.B., et al., Fluorescence-activated cell sorting and phenotypic characterization of human fibro-adipogenic progenitors. STAR Protoc, 2023. 4(1): p. 102008.

37. Patro, R., et al., Salmon provides fast and bias-aware quantification of transcript expression. Nat Methods, 2017. 14(4): p. 417–419.

38. Thul, P.J. and C. Lindskog, The human protein atlas: A spatial map of the human proteome. Protein Sci, 2018. 27(1): p. 233–244.

39. Love, M.I., W. Huber, and S. Anders, Moderated estimation of fold change and dispersion for RNA-seq data with DESeq2. Genome Biol, 2014. 15(12): p. 550.

40. Andersen, O.E., et al., Ketone Monoester Increases Skeletal Muscle Power and Energy Turnover in Older but Not Young Men Without Affecting Metabolic Economy: A Controlled, Double Blind, Cross-Over Trial. Acta Physiol (Oxf), 2026. 242(3): p. e70161.

41. de Morree, A. and T.A. Rando, Regulation of adult stem cell quiescence and its functions in the maintenance of tissue integrity. Nat Rev Mol Cell Biol, 2023.

42. Fry, C.S., et al., Myogenic Progenitor Cells Control Extracellular Matrix Production by Fibroblasts during Skeletal Muscle Hypertrophy. Cell Stem Cell, 2017. 20(1): p. 56–69.

43. Brightwell, C.R., et al., A glitch in the matrix: the pivotal role for extracellular matrix remodeling during muscle hypertrophy. Am J Physiol Cell Physiol, 2022. 323(3): p. C763–C771.

44. Biferali, B., et al., Fibro-Adipogenic Progenitors Cross-Talk in Skeletal Muscle: The Social Network. Front Physiol, 2019. 10: p. 1074.

45. Uezumi, A., et al., Mesenchymal Bmp3b expression maintains skeletal muscle integrity and decreases in age-related sarcopenia. J Clin Invest, 2021. 131(1).

46. Lee, S.-J., Genetic Analysis of the Role of Proteolysis in the Activation of Latent Myostatin. PLoS ONE, 2008. 3(2): p. e1628.

47. Chen, J.L., et al., Specific targeting of TGF-β family ligands demonstrates distinct roles in the regulation of muscle mass in health and disease. Proceedings of the National Academy of Sciences, 2017. 114(26): p. 201620013.

48. Cui, C.Y., et al., Skewed macrophage polarization in aging skeletal muscle. Aging Cell, 2019. 18(6): p. e13032.

49. Moratal, C., et al., IL-1β- and IL-4-polarized macrophages have opposite effects on adipogenesis of intramuscular fibro-adipogenic progenitors in humans. Scientific Reports, 2018. 8(1).

50. Nakazeki, F., et al., Loss of periostin ameliorates adipose tissue inflammation and fibrosis in vivo. Scientific Reports, 2018. 8(1).

51. Wang, L., et al., A single-cell atlas of bovine skeletal muscle reveals mechanisms regulating intramuscular adipogenesis and fibrogenesis. Journal of Cachexia, Sarcopenia and Muscle, 2023. 14(5): p. 2152–2167.

52. Farup, J., et al., Pericyte response to contraction mode-specific resistance exercise training in human skeletal muscle. J Appl Physiol (1985), 2015. 119(10): p. 1053–63.

53. Godwin, J.S., et al., Extracellular matrix content and remodeling markers do not differ in college-aged men classified as higher- and lower-responders to resistance training. J Appl Physiol (1985), 2023.

54. Kurosawa, T., et al., Transgenic Expression of Bmp3b in Mesenchymal Progenitors Mitigates Age-Related Muscle Mass Loss and Neuromuscular Junction Degeneration. Int J Mol Sci, 2021. 22(19).

55. Tabula Muris, C., et al., Single-cell transcriptomics of 20 mouse organs creates a Tabula Muris. Nature, 2018. 562(7727): p. 367–372.

56. Sieljacks, P., et al., Muscle damage and repeated bout effect following blood flow restricted exercise. Eur J Appl Physiol, 2016. 116(3): p. 513–25.

57. Farup, J., et al., Blood flow restricted and traditional resistance training performed to fatigue produce equal muscle hypertrophy. Scand J Med Sci Sports, 2015. 25(6): p. 754–63.

58. Sorensen, J.R., et al., An altered response in macrophage phenotype following damage in aged human skeletal muscle: implications for skeletal muscle repair. FASEB J, 2019. 33(9): p. 10353–10368.

59. Roth, S.M., et al., Skeletal muscle satellite cell characteristics in young and older men and women after heavy resistance strength training. The journals of gerontology.Series A, Biological sciences and medical sciences, 2001. 56(6): p. B240–7.

60. Rodgers, J.T., et al., mTORC1 controls the adaptive transition of quiescent stem cells from G0 to G(Alert). Nature, 2014. 510(7505): p. 393–6.

61. Nederveen, J.P., et al., Altered muscle satellite cell activation following 16 wk of resistance training in young men. Am J Physiol Regul Integr Comp Physiol, 2017. 312(1): p. R85–r92.

62. Snijders, T., et al., Prolonged exercise training improves the acute type II muscle fibre satellite cell response in healthy older men. J Physiol, 2019. 597(1): p. 105–119.

63. Scott, B.R., et al., Exercise with blood flow restriction: an updated evidence-based approach for enhanced muscular development. Sports Med, 2015. 45(3): p. 313–25.

